# Modelling the Impact of Dominant Transport Pathways on Antarctic Krill Fishing Activity in the Southern Ocean

**DOI:** 10.1101/2025.02.12.637831

**Authors:** Cian Kelly, Ingrid Ellingsen, Ragnhild Daae, Morten Omholt Alver

## Abstract

Antarctic krill (*Euphasia superba*) are a key component in the Southern Ocean ecosystem, especially in the Atlantic sector, where the majority of the population is concentrated. The Norwegian commercial krill fishery exclusively targets three subareas in the Antarctic: the western Antarctic Peninsula, and the northern shelves of both the South Orkney Islands and South Georgia. Given it’s reliance on oceanic transport from other regions and the potential impact of rising sea temperatures on the northern habitat, the South Georgian krill population is particularly sensitive to altering environmental conditions. The relative distance from the peninsular regions to South Georgia means that choosing to trawl in this region implies a higher risk, which is why it is exclusively targeted in winter when extensive sea-ice makes peninsular regions unsafe and inaccessible to commercial fishery operations. In this article, we show that relative to operations at South Orkney and the Antarctic Pensinsula, average catches at South Georgia have been lower with higher variability over the past 15 years. Using a Lagrangian modelling approach, we illustrate that variability in advection from source regions in the Antarctic Peninsula are correlated with proceeding catch values at South Georgia. This was not the case for source release sites at the South Orkney Islands. The dominant transport pathways for krill were strongly determined by position of regional fronts and the source sites of recruits to South Georgia were related to the position of fronts at both the Antarctic Peninsula and South Orkney Islands. This study highlights the importance of advective patterns on the variability in krill fishing activity and supports the hypothesis that South Georgia is a sink region for krill in the Southern Ocean while the western Antarctic Peninsula is a central source site.

## Introduction

Antarctic krill (*Euphasia superba*), hereafter krill, play a central role in the Southern Ocean ecosystem, where they form a crucial link between phytoplankton and higher trophic levels. Given they form dense swarms, migrate vertically, forage and graze at high rates and deposit fast sinking detrital pellets, they contribute greatly to carbon transport and sequestration [1]. It has recently been estimated that Antarctic krill faecal pellets sequester 20 MtC per productive season (spring to early autumn) [2]. Additionally, krill biomass is among the greatest of any marine species, with one of the largest zooplankton fisheries making it a key species for research and management.

Although the potential habitat for krill is quite large, they exhibit a highly heterogenous spatial distribution, with relatively higher biomass in the Atlantic sector estimated at 62.6 megatonnes [3, 4]. It is estimated 70% of the total stock is concentrated between longitudes 0° and 90°W [5]. Furthermore, the habitat in the Atlantic sector is partitioned between different life stages, with adult krill generally exhibiting a more oceanic distribution relative to earlier life stages [6]. This spatial heterogeneity is driven by many factors including but not limited to: the mortality in early life stages caused by salp blooms, the need for adequate bottom depth for the descent-ascent cycle of krill larvae, the sea-ice extent during austral winter, the temperature range in the northern extents of krill geographic distribution and trade-offs between predation risk and prey concentration [3, 5, 7, 8]. Most crucially, the dominant Antarctic Circumpolar Current (ACC) and its associated fronts are key to advecting krill away from centres of production and maintaining regional populations [3].

Fishing activity in the Atlantic sector (CCAMLR Area 48) reflects this spatial heterogeneity with the majority of activity taking place in three key subareas: the western Antarctic Peninsula (AP), the South Orkney islands (SO) and South Georgia island (SG), subareas 48.1, 48.2 and 48.3 respectively (Figure 1). The South Sandwich Islands, Subarea 48.4, are not targeted, likely due to harsh operating conditions in addition to the lack of shelf areas where krill swarms can aggregate in predictable hotspots for the fishery [9]. Catches in Area 48 have exceeded 400 thousand tonnes in recent years, with Norwegian fishing vessels harvesting a share of approximately 63% [10]. To monitor the condition of stocks in Area 48, annual surveys collect acoustic and trawl data (see for example [11]). Understanding connectivity between AP, SO and SG and potential seasonal and interannual variability can limit negative consequences of fishing on juveniles, and sub-adults, and simultaneously limit the carbon footprint of industry [10]. This is particularly relevant for the sustainable exploitation of the krill fishery at the high latitude (∼ 54*^◦^*) SG habitat [9].

**Fig 1.**
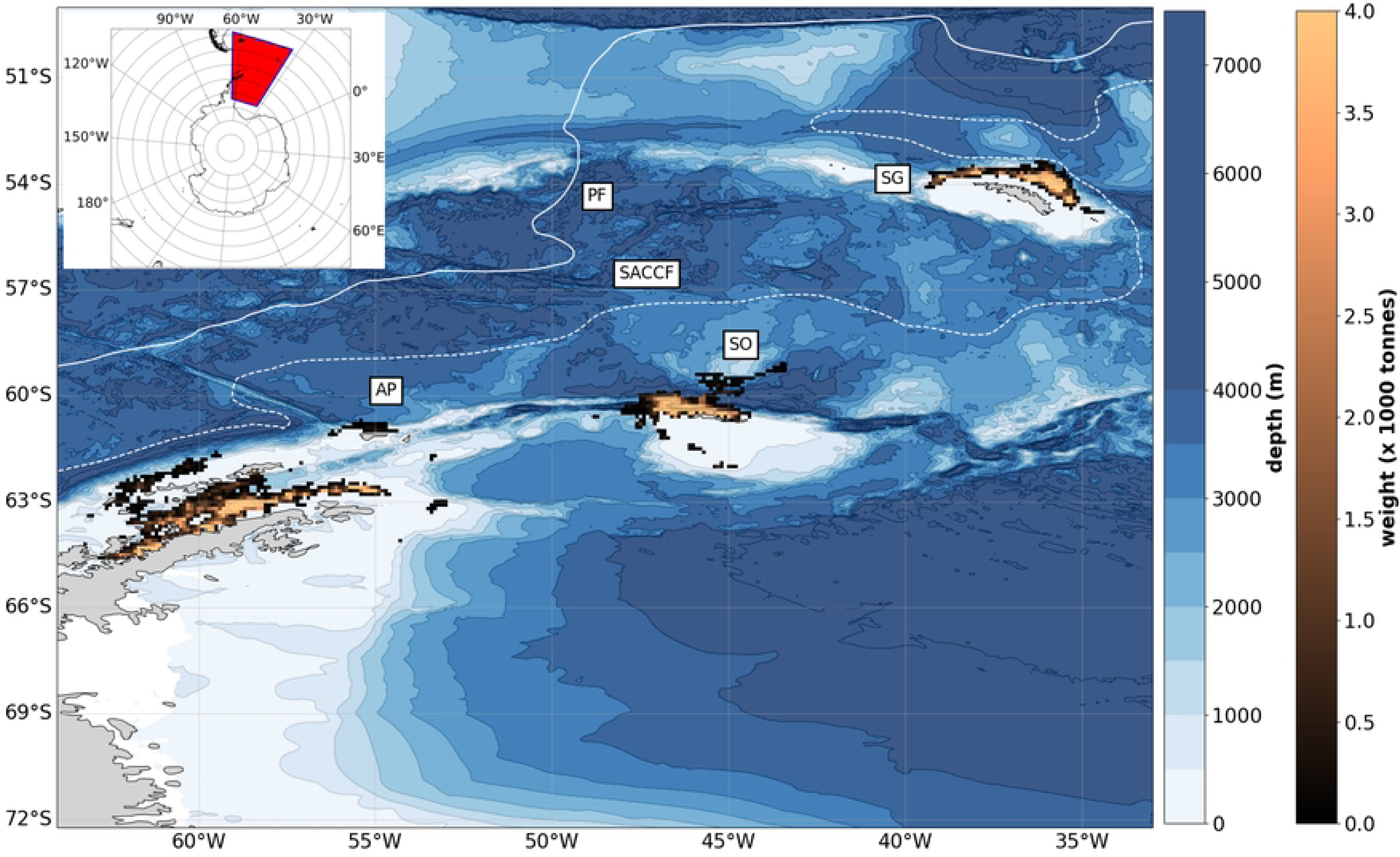
The copper colourmap shows the average catch values (x 1000 tonnes) of Antarctic krill by the Norwegian krill fishery discretised in 0.1◦×0.1◦ bins from 2006 to 2023 (CCAMLR data). The major fishing regions are annotated as follows: Antarctic Peninsula (AP), South Orkney Islands (SO) and South Georgia (SG), which correspond to data from CCAMLR subareas 48.1, 48.2 and 48.3 respectively. The solid and dotted white line show the mean position of the Polar Front (PF) and Southern ACC front (SACCF) respectively (from [24]). The blue colourmap shows the domain bathymetry from GEBCO with contours at 500m intervals from 0 to 8000m.

Modelling studies have elucidated the drivers of spatial connectivity between AP, SO and SG regions. SG is bounded by the southern ACC front (SACCF) to the south and the Polar Front (PF) to the north (Figure 1). The SACCF approaches SG from the southwest is the main transport pathway for krill to SG from the Anarctic Peninsula [12, 13]. A fraction of the biomass is also advected eastward to the South Sandwich Islands which is not targeted by the fishery. Krill in high biomass regions off SG in summer have been confirmed to originate from regions covered in sea-ice during winter [12]. In addition, acoustic studies have found periodic fluctuations in krill biomass at SG every 4-5 years, associated with preceding August SST [14]. Finally, it is postulated there will be a southward shift in krill biomass, which would heavily impact the local SG population [15]. Given the SG krill population is not considered a self-sustaining population and therefore relies on transport into the region, oceanographic conditions such as prevailing current directions, transient eddy formations and the position of oceanic fronts are likely important in recruitment to the SG shelf. Although there are numerous studies that have investigated transport to SG, there are none that have compared these patterns to associated Norwegian fishing activity (the major krill fishery in Area 48).

The distances between the southern and northern extent of the krill fishery are vast (∼ 10*^◦^* latitude range, see Figure 1) and thus decisions on harvesting locations are central to sustainable expoitation. There are many factors that influence the decision of fishing vessels on when and where to trawl including weather conditions, quotas and auction prices, and the intensity of activity is not simply a function of target species distribution [16]. Regardless, we hypothesize that variability in advective patterns will have a large influence on the average catch year on year. In this study, we explore the relative impact of recruitment patterns from AP and SO on subsequent fishing activity at SG. Firstly, we explore the seasonal and interannual variability in catch data from Norwegian fisheries in the area to deduce the trends in the past decade and a half.

Secondly, we detail a Lagrangian modelling approach used to analyse transport between AP, SO and SG, both identifying origins of and transport to SG from AP and SO predicted by model simulations over a 15 year time period. Finally, we analyse these model statistics in relation to trawl data to infer correlations between modelled transport patterns and interannual variability in fishing activity in the studied regions.

## Materials and methods

### Trawl data

Trawling data (data product 680) from the Norwegian krill fishery in the Southern Ocean was granted on request from CCAMLR as of the 15th of April 2024, with data covering the period from the January 1st 2006 to December 31st 2023 selected for analysis. The data consisted of 140,760 disaggregated observations from three Norwegian fishing vessels operating in ubareas 48.1 (western Antarctic Peninsula region), 48.2 (north of South Orkney Islands), and 48.3 (northern shelf of South Georgia), referred to hereafter as AP, SO, and SG, respectively. Each observation recorded an individual krill trawl recorded by one of the fishing vessels.

The variables included in the analysis were krill catch weight (tonnes), latitude and longitude coordinates of the trawl initiation, and the time of the trawl initiation. The spatial extent of the fishery (Figure 1) was studied in order to inform the initialisation parameters for Lagrangian simulations described in the following section and to compare against spatial probability fields predicted from the model (both origin and transport pathways of krill). In addition, seasonal and interannual trends in trawl data were investigated to illustrate and contrast variability in regional catch values.

### Lagrangian Particle Tracking

#### Model Setup

To simulate the particle model forward in time, global ocean physics model data was used as input. Model velocity fields from the global physics reanalysis GLORYS12V1 product, hosted on the Copernicus Marine Service, were acquired to force the Lagrangian particle tracking module [17]. This product is the output from NEMO ocean model simulations, which are atmospherically forced every 24 hours and utilize a reduced-order Kalman filter for data assimilation. The model operates on a regular Arakawa C grid with a horizontal resolution of 1/12*^◦^* (approximately 8 km, depending on latitude) and 50 vertical layers.

In this analysis, we extracted horizontal u- and v-components of seawater velocity from the daily dataset. Model data from 2006 to 2020 inclusive were downloaded in netCDF format using the *copernicusmarine* Python API. This period corresponds to the overlap in available model and observational data. The horizontal u- and v-components of seawater velocity were used as input to *OpenDrift* in offline simulations. *OpenDrift* is an open-source framework for Lagrangian particle modelling scripted in Python, which allows flexibility in model configuration for various applications [18].

The transport of passive particles was simulated using a second-order Runge-Kutta propagation scheme. Particles were deactivated if they interacted with the coastline and were not included in subsequent analysis. Output data were saved in netCDF format according to CF conventions for the storage of trajectory data.

#### Simulation and Analysis of Particle Transport

Simulations were run with computational resources from SAGA, a supercomputer which is part of the *Sigma2* national infrastructure in Norway. Particles (n = 1 x 10^4^) were released from AP and SO at 75m depth, a high mean density krill layer, in a regular rectangular grid with equal spacing, every 7 days from August 1st each year for a total of 20 releases in each grid cell. This meant a total of 2 x 10^5^ particle releases per each of the 15 simulation years. The time step of 10 minutes was chosen to sufficiently sample the model fields with a model time period of 300 days to allow particles released in the preceding year, with the earliest in August, to reach SG by austral winter the following year.

The AP scenario spanned a meridional range from 63*^◦^*W west of Antwerp Island to 54*^◦^*W at Clarence Island, the easternmost island in the South Shetland Islands. Zonally, it spanned from 65*^◦^*S south of Antwerp Island to 60.5*^◦^*S at the shelf break north of Elephant Island. This coverage is analogous to CCAMLR Subarea 48.1, except for the omission of the eastern sector of the Antarctic Peninsula. The SO scenario spanned a meridional range from west from 48*^◦^*W west of Coronation Island to 44.35*^◦^*W east of Laurie Island. Zonally, it ranged from 61*^◦^*S south of the South Orkney Islands to 59.2*^◦^*S at the shelf break. This covers the western and northern SO shelf.

Lagrangian dynamics operate in a continuum and thus to analyse connectivity between predefined regions, it is convenient to discretise particle trajectory data (x and y coordinates in this case). We first performed an analysis to quantify the probability krill recruited to SG originated in sites from AP and SO. The total number of particles reaching SG from each release site was summed over the 20 releases inside 1*^◦^*× 1*^◦^* geographical bins. This calculation was averaged and normalised over the 15 modelled years in order to assign a relative probability of origin of krill arriving in SG from locations in AP and SO.

Similarly, we performed a unique occupancy calculation which counted the proportion unique particles visiting 0.1*^◦^*× 0.1*^◦^* grid cells, averaged across the 15 years of simulations and 20 release dates. Values could therefore range between 0 and 100%, although the probability of a particle visiting any individual grid cell is quite low. Dominant transport pathways of passive drifters are often deduced from this calculation (see for example [19, 20]).

Finally, we analysed the total percentage recruited and the transit times from AP and SO to SG across releases and years to investigate interannual and seasonal variability in recruitment respectively. Recruitment statistics were then compared to trawl data to infer correlations between modelled transport patterns and interannual trends in Norwegian fishing activity in the studied regions.

## Results

### Spatiotemporal variability in Norwegian fishing activity (2006 - 2023)

#### Spatial variability

Figure 1 illustrates clearly the localised effort of Norwegian fishing activity over the past two decades (2006-2023) both in relation to bathymetry of the continental shelf and the mean position of the SACCF. At AP the Bransfield and Gerlache Straits are the main focus of the fishery, with some activity north and west of the South Shetland Islands and north of Elephant Island. At SO, the fishery is concentrated northwest of Coronation Island over the northern shelf breaks which is a region with several glacially eroded troughs [21]. At SG the northern part of the island south of the SACCF at the shelf break is most targeted by the fishery. Frontal regions and circulation patterns north and specifically, northwest of the island, are known to be productive regions for krill [22, 23]. It is argued that CCAMLR managment policy should focus on shelf areas with elevated bathymetry, in tandem with considerations regarding regional climate change [9]. Results here demonstrate the importance of shelf areas to the fishery.

#### Catch variability

During the study period, the distribution of catch events across regions was as follows: 35% in AP, 41% in SO, and 24% in SG. The SO scenario exhibited the highest average catches with intermediate variation, while the AP scenario had intermediate average catches and the lowest variation. In contrast, the SG scenario recorded the lowest average catches but the highest variation (Table 1). This combination of low catch values and high variability, along with SG’s considerable distance from peninsular regions (approx. 1000–1500 km), suggests that SG represents a relatively unpredictable fishery. Additionally, the high coefficient of variation (CV) across all subareas highlights the inherent uncertainty in krill fisheries.

**Table 1.**
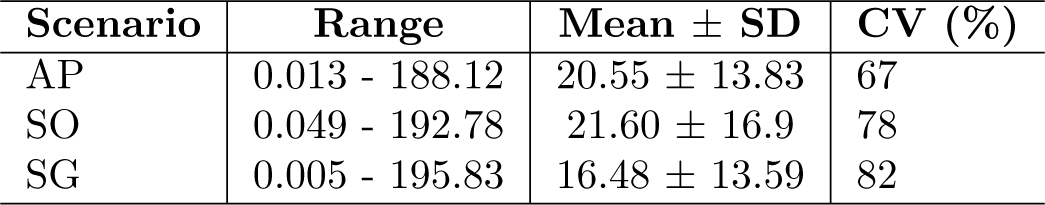
Summary of krill catch statistics in subareas (AP, SO, SG). The table presents the range of catches, mean catch values with standard deviation (Mean ± SD), and the coefficient of variation (CV) expressed as a percentage.

| Scenario | Range | Mean ± SD | CV (%) |
| --- | --- | --- | --- |
| AP | 0.013 - 188.12 | 20.55 ± 13.83 | 67 |
| SO | 0.049 - 192.78 | 21.60 ± 16.9 | 78 |
| SG | 0.005 - 195.83 | 16.48 ± 13.59 | 82 |

#### Temporal variability

Norwegian fishing activity is seasonally distinct, where SO is targeted in austral summer, AP in autumn and spring, while SG is targeting in austral winter (Figure 2 a).

**Fig 2.**
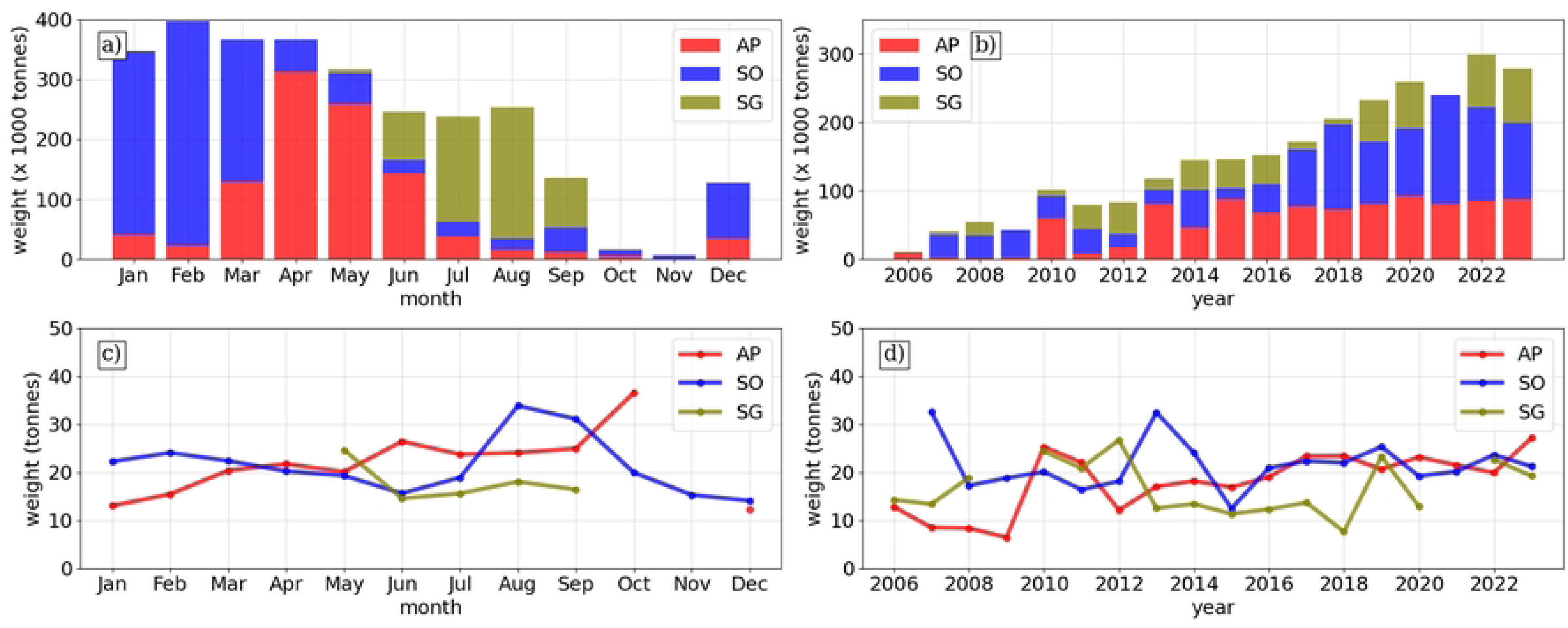
Monthly and yearly distribution of Norwegian catch events from three fishing areas in the Southern Ocean (AP: Antarctic Peninsula; SO: South Orkney Islands; SG: South Georgia)

Austral summer is the most active period while fishing turns to SG in winter when sea-ice makes peninsular regions inaccessible. In July and August fishing is almost exclusively at SG during the period studied. There has been an increase in targeting of krill over the past two decades, with catches rising above 200 kilotonnes in 2018 from annual yields below 100 kilotonnes in the early 2000s (Figure 2 b).

The average catch appears to be higher in months with less total activity (Figure 2 c). This could indicate fishers knowledge of favourable conditions outside the main fishing season (take for example SO in August, AP in October and SG in May).

Otherwise, August and September have particularly high catches, which could support the hypothesis that colder conditions are favourable for the fishery, where September exhibits the lowest SST on average. The relatively low average catches at SG suggests a challenge with finding swarms of equivalent density to those found at AP and SO (Figure 2 d). Also, since 2016, AP and SO average catches appear more stable (∼25 tonnes), whereas SG average catches have fluctuated between 10-23 tonnes.

### Connectivity between major krill fishing regions

#### Particle trajectories

Figure 3 shows snapshots of the Lagrangian model over 10 months from release sites at AP and SO regions in October 2019 until the majority of subsetted particles have reached SG in July 2020, a period of concentrated fishing activity. Particles from AP are coupled strongly to the SACCF where they first are advected into regions north of SO, spending time in the SO shelf area before following the SACCF north to enter SG. In contrast, SO particles, advected east and southeast before topographic steering of the currents drives particles north and up onto the SG shelf. The general transport patterns from SO are in line with previous studies [19].

**Fig 3.**
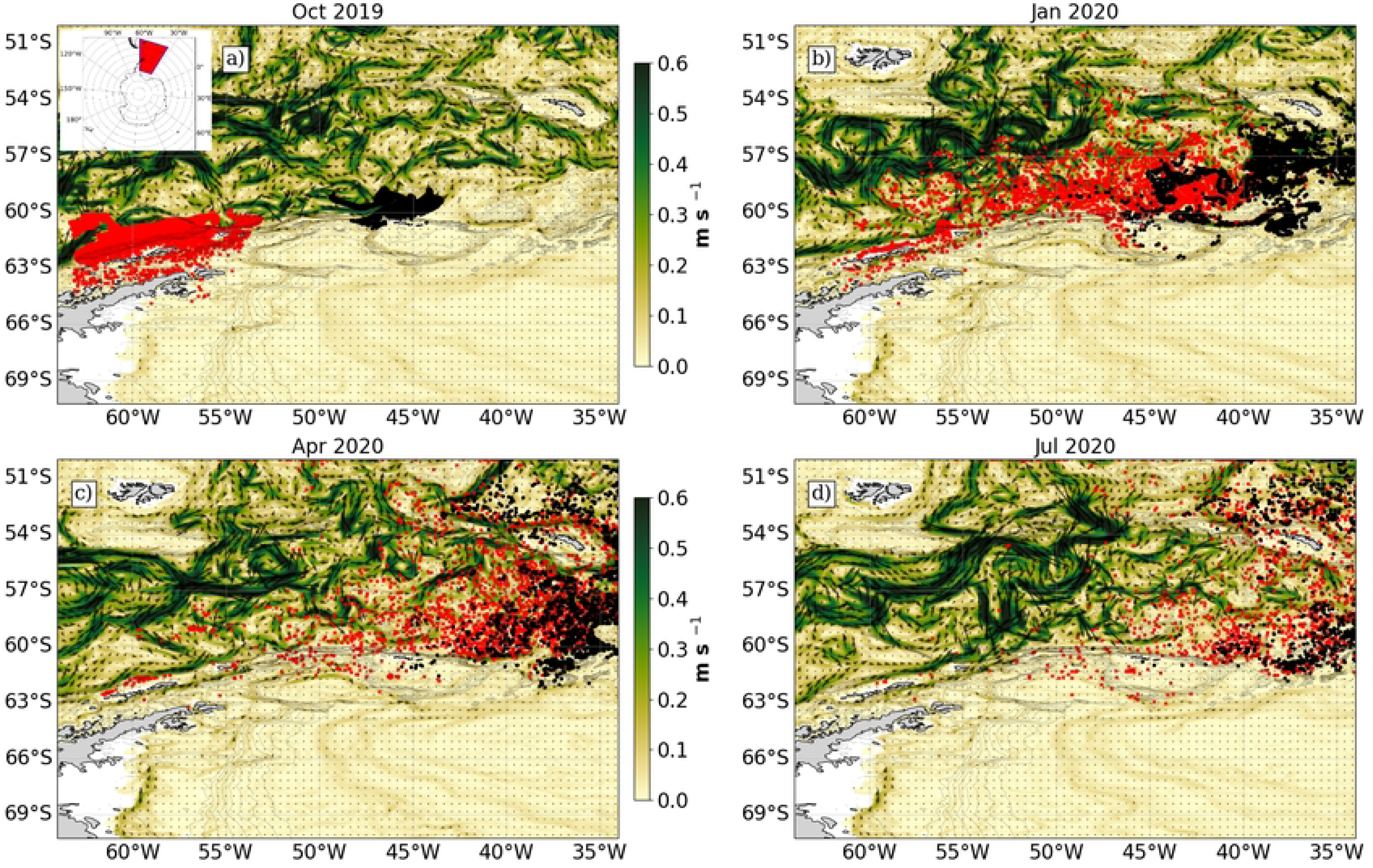
Snapshots of the Lagrangian model at 4 time stamps for particles released in October 2019 (a: October 2019, b: January 2020, c: April 2020, d: July 2020) with the quiver map indicating the zonal and meridional direction of currents and the colourmap showing the total magnitude of the horizontal surface currents (u- and v-components). The red and white points illustrate the paths of particles released from Antarctic Peninsula (AP) and South Orkney Islands (SO) sites respectively.

#### Particle origins

The relative probability of krill origins were analysed to identify major subarea sources of krill to SG and overlap with krill fisheries for the period simulated. In the case of AP (Figure 4 a), the highest probability values were from areas overlapping with SACCF, with areas north of the South Shetland Islands and Elephant Island the most important source areas, where over 70% of SG krill originated. This overlaps with major fishing regions. Inversely, the Bransfield Strait and Gerlache Straits were less important source areas, although they are intensely harvested (Figure 1). At SO, the probability map followed a T-shaped spatial distribution with bank area region north of the the shelf at approx ∼60*^◦^*S spatially overlapping with high fishing activity (Figure 4 b). There was less overlap northwest of the island close to the shelf, where the majority of the fishing activity takes place. The origin of krill to SG was localised to the shelf breaks in both scenarios, which are known to be highly productive regions.

**Fig 4.**
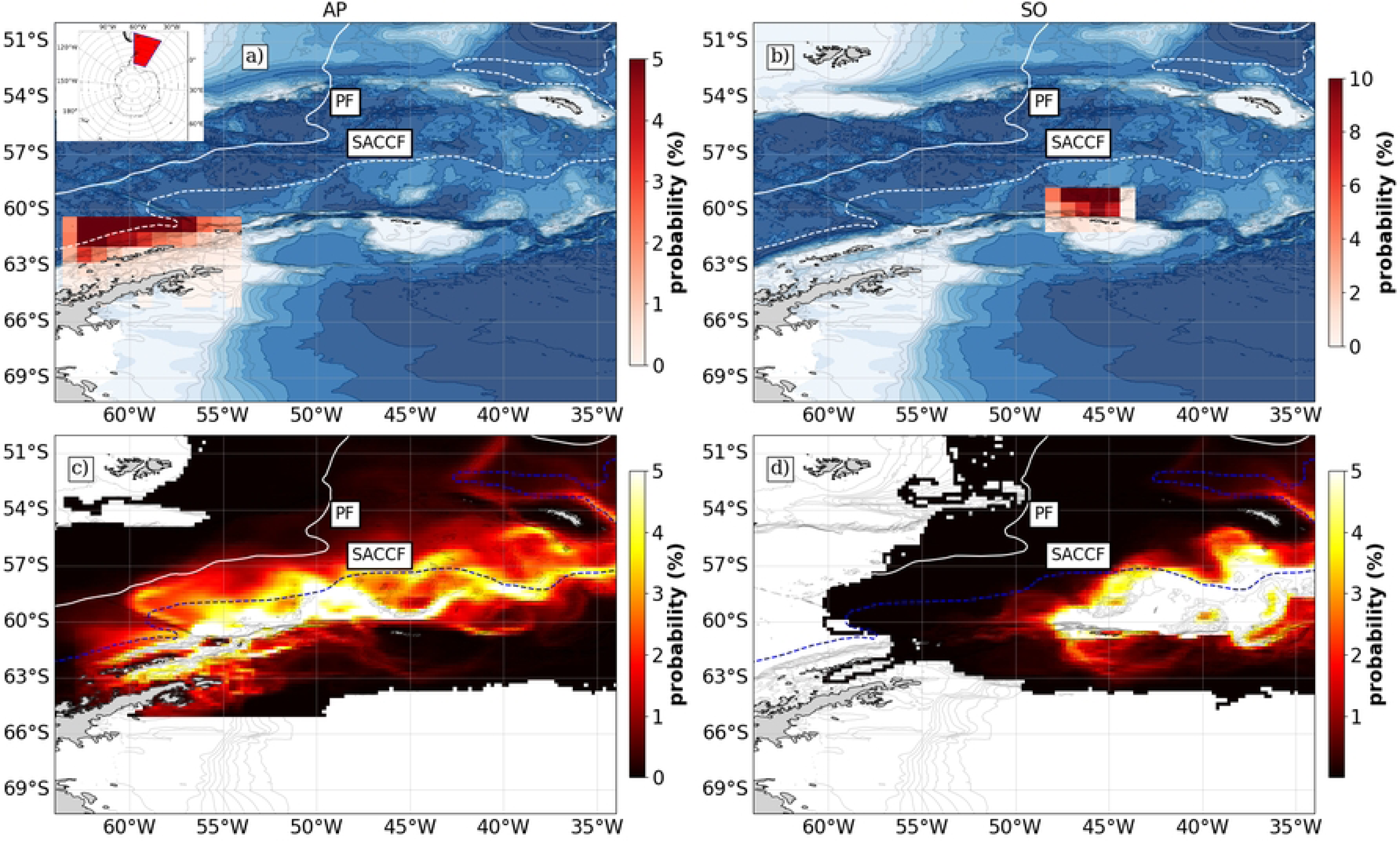
Average probability maps across all simulations (15 years × 20 releases). Panels a) and b) show the relative probability of origin of particles in discrete 1*^◦^*× 1*^◦^* bins from source sites at Antarctic Peninsula and South Orkney Islands. In contrast, panels c) and d) show the absolute probability a particle will occupy a 0.1*^◦^*× 0.1*^◦^* grid cell during the model simulation.

#### Dominant pathways

The unique occupancy plots across all simulations (15 years × 20 releases) show the general trends in dominant pathways for all particles across all simulations give a picture of the total advective patterns (Figure 4 c and d). From the initial regions, particles follow two main branches from AP, the through the Bransfield Strait and north of the South Shetland Islands. Some transport is north towards the PF which acts as a barrier to further transport (Figure 4 c).

From 45*^◦^*W, north of SO is a choke region for krill transport, with particles either following the SACCF north and approaching SG from southwest of the island, or approaching the island from the southeast and arriving on the northern shelf where they enter regions accessible by fishers. From here, they are either retained in on the northern part of the shelf or follow the SACCF and transit northeast. From SO there is a greater dispersal of particles from the source region (Figure 4 d).

Many particles are transported east before a northbound transit to SG, but much of the krill population is also transported east to the South Sandwich Islands, an area with no fishing activity and there is quite limited knowledge on krill distribution.

### Seasonal and interannual variability in South Georgia recruitment patterns

#### Seasonal variability

Both the percentage recruited and the time to reach SG from AP was stable with low variation around the interannual mean (Figure 5 a). Mean values fell in the range of 7-10%, a quite high figure. There is a slight uptick in the fraction recruited from October to December which may relate to receding sea-ice around the peninsular region in this period. Sea-ice can reduce the zonal transport of krill across the Southern Ocean [13]. In comparison, there was higher variability in the number of recruits from SO around the interannual mean, but no clear seasonal pattern (Figure 5 b). The percentage recruited was lower ranging with mean values between 4-6%, although there are individual years where recruitment peaked at 10% showing the volatility in recruitment patterns from SO.

**Fig 5.**
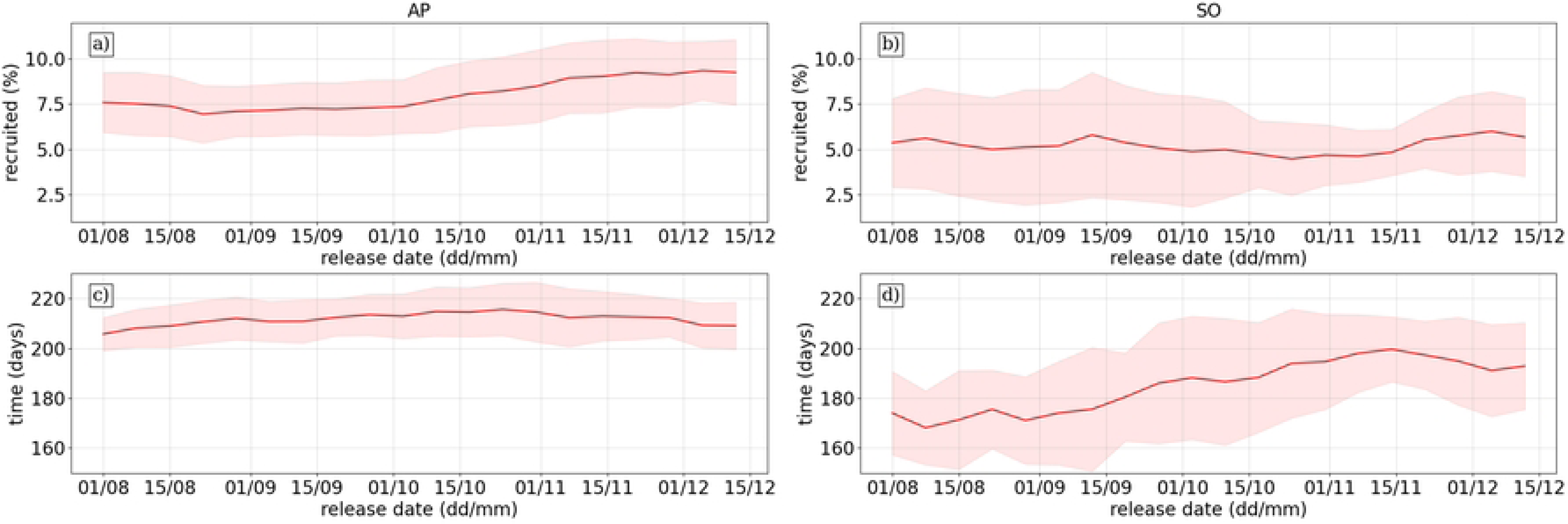
Recruitment statistics over release dates to South Georgia from source regions, where each release date from source site is represented by the interannual mean (red line) and standard deviation across the 15 simulation years (red shaded region). The top row shows the average percentage recruited to South Georgia from AP (a) and SO (b), while the bottom row shows the time taken for the average time in days to reach South Georgia from AP (c) and SO (d).

The time taken to reach SG from AP is relatively stable, ranging from between 200 to 220 days on average, with little interannual variation (Figure 5 c). This may reflect stability in the mean ACC flow speeds associated with the SACCF. Again, there is greater variability in the SO simulations, where average recruitment time can range from between 170 to 200 days, with high variation among the simulated years (Figure 5 d). This reflects the wide spread in mean transport patterns from the dominant pathways analysis (Figure 4 d) Many particles released from SO took a more eastward path and looped anticyclonically around SG to enter the northern shelf, while a more direct path from the southern shelf is also prominent in the results.

#### Interannual variability

The frequency histograms of average release statistics reflect these patterns showing a wider spread in both time and percentage recruited from SO relative to AP releases. It also shows that the patterns follow a tight normal distribution for AP releases, centred on 215 transit days and 7% recruitment (Figure 6 a and c), illustrating a predictability in the transport patterns which may be relevant to connectivity between AP and eastern regions. SO release statistics follow a lognormal distribution for percentage recruited, centred around 3-4% with a long tail from 5-14% (Figure 6 d) and a more normal distribution for transit times centred around 190 days (Figure 6 b).

**Fig 6.**
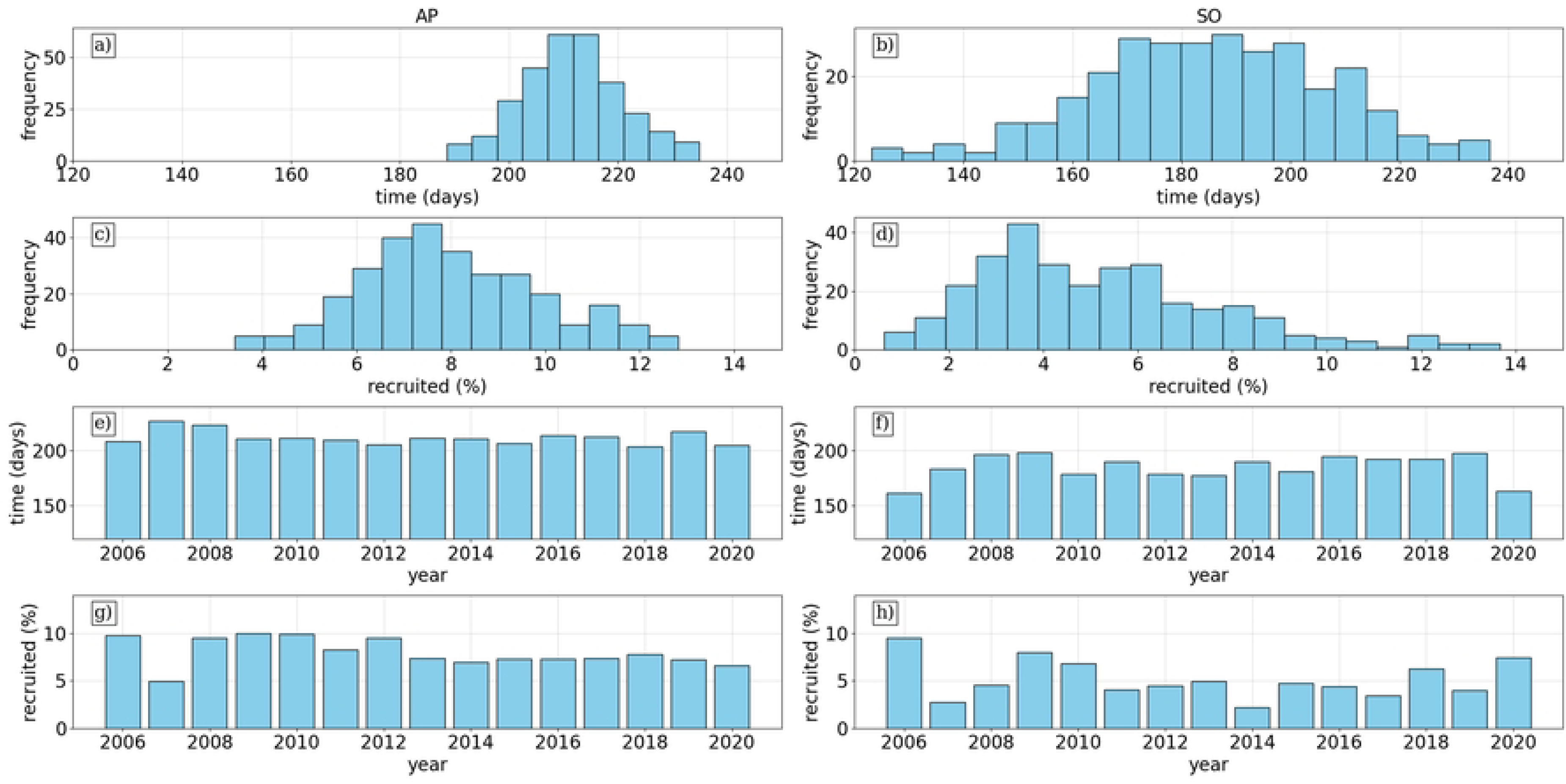
[a-d] Histograms of recruitment statistics averaged across releases for all years (15y x 20r simulations), where shade of blue reflects the relative frequency of the values. The first two rows show the time (days) to reach SG from AP (a) and SO (b) and percentage recruited from AP (c) and SO (d). [e-h] Histograms of statistics displayed across the years simulated with each bar representing the average recruited across 20 releases for time (days) from AP (e) and SO (f) and recruited percentage from AP (g) and SO (h).

Similarly, the interannual distribution of recruitment time is quite stable for AP (Figure 6 e) at just over 200 days in each simulation, while there are years where it takes on average 200 days but others closer to 150 days from SO (Figure 6 f). Finally, the percentage recruited does show some variability with highs close to 10 percent recruitment 2006, 2008, 2011 for example from AP (Figure 6 e), while SO has high variability with highest 2006, lows 2007 and 2014 in low single digits (Figure 6 f).

#### Correlations Between Recruitment and Fishing Activity

We examined the linear relationship between model and observation datasets by comparing annual averages to determine if interannual trends in fishing activity at SG correlate with recruitment statistics (Figure 7). Given the variability between individual fishing events, this approach helps to identify broader trends.

**Fig 7.**
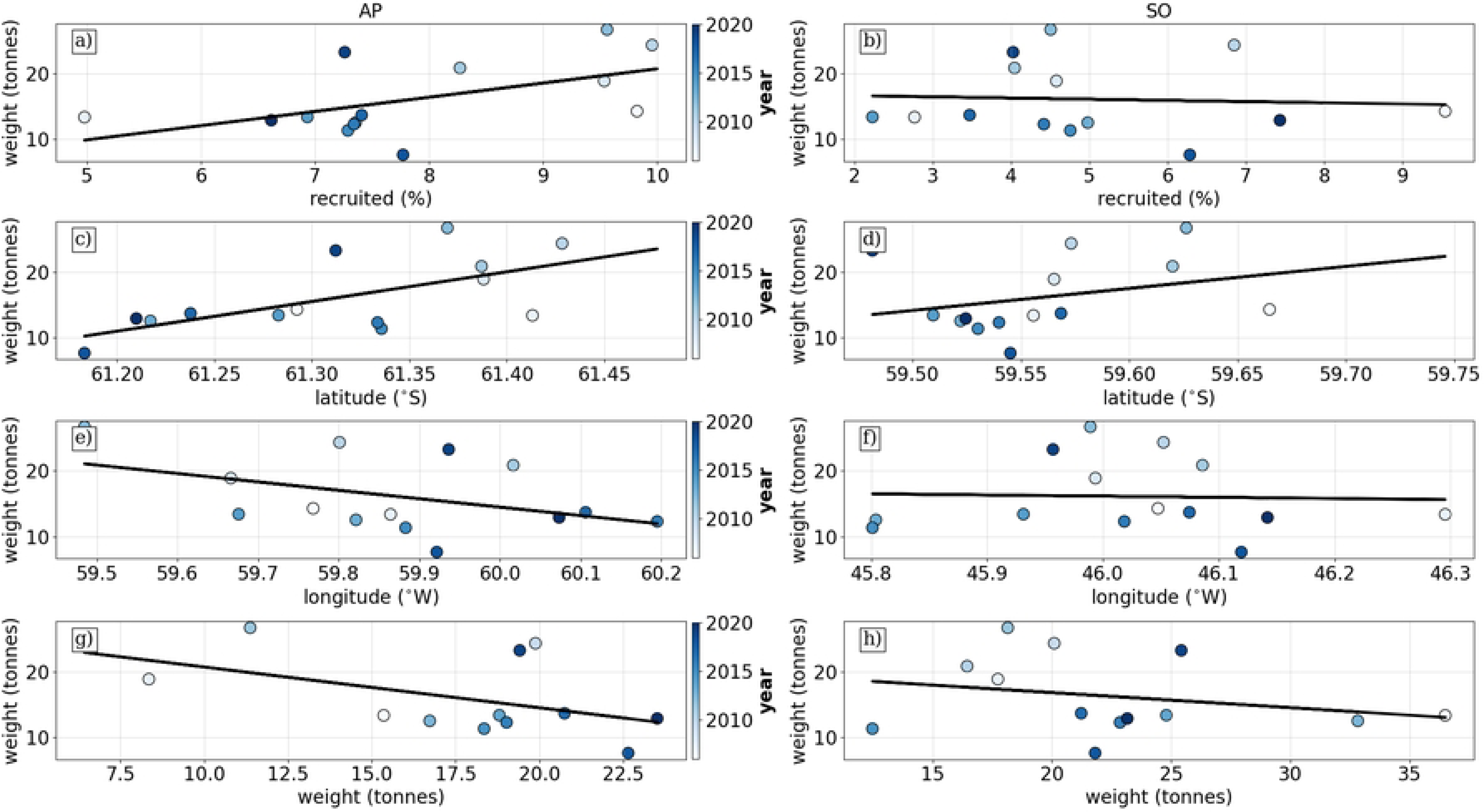
Linear relationships between Lagrangian transport model statistics and fishery catch data. The colourmap for points indicates the year. The top row shows the relationship between the percentage recruited to SG and the average catch at SG from a) AP and b) SO. Rows 2-3 show the relationship between the latitude and longitude centre of gravity of origins of SG krill from AP (c and e) and SO (d and f). Finally, we show the relationship between the average preceeding catches at AP (g) and SO (h) compared to average proceeding catches at SG.

For the AP scenario, the correlation between the percentage recruited to SG and fishing events was significant (see Table 2), indicating that recruitment from AP source sites significantly influences SG catch. In contrast, the correlation for SO percentage recruited was not significant, suggesting a lesser impact from SO sources.

**Table 2.** Linear Regression Results for AP and SO scenarios where the Response Variable was the average catch at SG and Predictor variables were: percentage recruited from source, latitude and longitude centre of gravity of source particles and weight of catch at source site prior to SG fishing season.

| Scenario | Predictor | $\beta_0$ | $\beta_1$ | SE | $R^2$ | p-value |
| --- | --- | --- | --- | --- | --- | --- |
| AP | recruited (%) | -0.897 | 2.164 | 0.969 | 0.294 | **0.045** |
|  | latitude (°S) | -2755.868 | 45.210 | 15.917 | 0.402 | **0.015** |
|  | longitude (°W) | 784.119 | -12.827 | 7.708 | 0.187 | 0.122 |
|  | weight (tonnes) | 26.952 | -0.620 | 0.382 | 0.209 | 0.135 |
| SO | recruited (%) | 17.025 | -0.181 | 0.838 | 0.004 | 0.833 |
|  | latitude (°S) | -1982.76 | 33.561 | 31.444 | 0.087 | 0.307 |
|  | longitude (°W) | 97.291 | -1.764 | 12.638 | 0.002 | 0.891 |
|  | weight (tonnes) | 21.437 | -0.229 | 0.265 | 0.064 | 0.405 |

The Centre of Gravity (CG) of latitude for AP origins to SG showed a significant relationship (see Table 2), while the longitude relationship was weaker. For SO origins, both latitude and longitude exhibited much weaker relationships.

Finally, there were no clear relationships between preceding catches at AP or SO, indicating that catch levels at SG are not predicted by earlier catches at other locations.

## Discussion

### Variability in fishing activity

#### Trends in fishing activity

Results from the analysis of Norwegian krill fishing activity from 2009-2023 indicate a clear localisation of fishing activity to shelf areas in the western Antarctic Peninsula (AP), north of South Orkney (SO) and South Georgia (SG), with activity bounded by the Southern Antarctic Circumpolar Current Front (SACCF) to the south and the Polar Front (PF) further north (Figure 1). All activity was recorded in the Atlantic sector for this period in CCAMLR subareas 48.1, 48.2, and 48.3, illustrating the density of effort in recurring locations. Fishing activity has increased over the period investigated, putting relatively higher pressure on local krill populations. Catches have increased from below 100 kilotonnes annually from 2006-2012, to over 250 kilotonnes in 2020, 2021, and 2023.

#### Impact of Environmental Factors and Future Considerations

This increase in fishing activity is particularly relevant for SG, where seawater temperature is often above the thermal limit of the krill. Increases in temperature will likely cause contractions in the habitat further south [15]. SG represents the northern extent (53°S) of both fishing activity and krill habitat and is somewhat disconnected from the other 75% of recorded catches south of 60° latitude (AP and SO) (Figure 1). Our results show higher variability in fishing activity at SG relative to other regions, both in terms of average and absolute catch (Figure 2 b) and overall CV of 82% across all recorded catches). As SG is a winter fishery, targeting proceeds in SO and AP when sea-ice makes these regions inaccessible. Fishers may choose to omit operations in years where yields are particularly high at SO or AP (e.g., 2021 where recorded catches for AP and SO inclusive approached 250 kilotonnes). The SG fishery is subject to volatility in yields and, while it is commercially viable to target the region now, it may not be 5-10 years from now given current rates of warming [15]. To operate in uncertain conditions may require adaptation of the fishery. For example, fishing companies are increasingly considering using data sources for planning and routing, developing sophisticated decision support systems which can allow more targeted, and economically and environmentally sustainable fishing activity [16].

### Importance of Antarctic Peninsula as source region for South Georgia

#### Transport pathways

Particles released from AP and SO sites that reached SG followed a northeastern path following the ACC and it’s associated fronts. Around nine months later, a significant proportion entered the South Georgia shelf from a southeastern entry point. 3. This transport pathway is influenced by the proximity of the SACCF to the shelf edge in western AP, where strong incursions of Circumpolar Deep Water enhance productivity, making it an important krill source region in the Atlantic sector [13]. Our results further corroborate previous findings, showing that particles entrain in currents near the Bransfield Strait and between the shelf edge and the South Shetland Islands (Figure 4c). The highest recruitment to SG originates from the shelf edge north of the South Shetland Islands and Elephant Island (Figure 4a), highlighting key regions of krill supply.

#### Overlap with fishing activity

The observed transport pathways of krill influence their availability to fisheries, with key differences between AP and SO. In AP, there is strong transport within the Bransfield and Gerlache straits, key regions for the fishery (Figure 1). There is also significant overlap in fishing activity and sources of SG krill over the 15 years simulated, particularly at the bank north of the island (Figure 4b). The distinct transport patterns from AP and SO are shaped by regional oceanographic features. A critical chokepoint for krill transport occurs between 50*^◦^*W to 45*^◦^*W where strong flows associated with the Weddell Front constrain AP-derived krill to distinct pathways (Figure 4 d). This leads to a more concentrated recruitment pattern at SG compared to the more dispersive transport from SO release points. The northeastward route taken by particles from sites east of 40*^◦^*W aligns with previous studies [19], reinforcing the role of ocean currents in influencing krill availability to the SG fishery.

#### Recruitment statistics

The differences in recruitment statistics between AP and SO reflect underlying variations in transport dynamics. On average, 8% of particles released from AP were recruited to SG 6 a) compared to 5% from SO 6 b). Despite this lower recruitment rate, SO particles reached SG earlier, with a mean transport time of 190 days, compared to 210 days from AP. The greater spread in SO recruitment times suggests higher variability in transport patterns, likely due to more dispersive currents influencing SO pathways. In contrast, AP-derived krill experience relatively stable transport, with stronger confinement to well-defined pathways, leading to more consistent recruitment rates. This is further supported by previous modeling efforts which found a higher recruitment fraction from SO to SG with a shorter mean transport time when using the OCCAM velocity field (0.25° × 0.25° resolution). The difference in transport times between studies suggests that the resolution of the velocity field may influence recruitment estimates. Notably, the mean transport time from AP to SG in [12] was 160 days, which is less than all 15 year x 20 release simulations conducted on the higher-resolution (0.08°) grid, potentially indicating that finer-scale processes impact transport duration. For example, north of SO is known for strong sub-mesoscale and mesoscale eddy activity which may influence such calculations.

#### Interannual variability

The differences in interannual variability between AP and SO transport pathways suggest distinct underlying oceanographic drivers. The SO simulation exhibited a larger spread in both release dates and interannual variability, whereas AP releases showed lower relative variability. This pattern likely reflects the relative stability of the zonal current velocities from AP to SG, which provide more consistent transport conditions. In contrast, SO-to-SG transport is influenced by more variable mesoscale and submesoscale processes, leading to greater year-to-year fluctuations.

Despite the lower overall variability in AP transport, interannual changes in AP releases had a measurable impact on average catches at SG. Specifically, fluctuations in recruitment percentage and source location. (Figure 7) suggest that variations in AP transport can influence the availability of krill to the fishery. This implies that certain years may experience stronger recruitment pulses from AP, potentially affecting krill biomass at SG.

Conversely, recruitment from SO showed weak relationships with interannual variability, indicating that SO-derived krill play a less significant role in determining the krill available to the SG fishery in the following season. This may be due to the more dispersive transport pathways from SO, which reduce the direct impact of variability in source conditions on krill availability at SG. Additionally, the broader spread of SO particles may result in a more diluted and less predictable recruitment signal.

These findings highlight the importance of AP as a relatively stable and influential source of krill for SG, while SO’s contribution is more variable and less directly linked to interannual recruitment trends at SG. Understanding these dynamics is crucial for predicting krill availability and managing fisheries in a changing ocean environment.

## Limitations and Further Considerations

This study focused on passive drift to assess large-scale krill transport pathways but did not incorporate active krill behavior, which can significantly influence long-range and long-term advection. For instance, sea-ice-associated behavior plays some role in krill distribution by affecting feeding patterns and modifying transport dynamics [13].

Seasonal sea-ice expansion and retreat can alter krill retention times in specific regions, potentially impacting recruitment to South Georgia (SG).

Diel vertical migration (DVM), a primary behavioral trait of krill, also affects transport outcomes. Krill migrate to surface waters at night and descend to deeper layers during the day, where current velocities and directions differ. This behavior can influence advection rates, as deeper currents often move at different speeds and directions than surface currents.

In addition to behavioral factors, physical oceanographic processes such as upwelling can impact krill availability at both source and sink regions. Upwelling brings nutrient-rich deep water to the surface, enhancing primary production and increasing food availability for krill populations. This process can influence krill survival rates at the Antarctic Peninsula (AP) and South Orkney (SO) source regions, as well as their growth and aggregation near SG. Variability in upwelling intensity could thus modulate krill biomass over interannual to decadal scales, affecting fisheries productivity.

A key physical constraint in krill transport is the presence of a chokepoint for particles moving onto the SG shelf, particularly around 45°W. Strong flows at this location regulate the influx of krill, potentially increasing biomass in the region and leading to spatial overlap with offshore fishing activity. Survey data imply similar historical patterns, indicating that the recruitment of krill to SG fisheries may be influenced by this transport bottleneck. Future studies should further investigate how regional hydrodynamics, including mesoscale eddies and wind-driven mixing, modify krill transport across this critical zone.

## Conclusion

The Antarctic Peninsula (AP) has experienced rapid warming since the 1950s [25], with significant implications for the South Georgia (SG) krill population. As a key source region for krill recruitment to SG, environmental changes in AP can have cascading effects on krill availability across the Atlantic sector. Given that SG represents the northernmost extent of krill habitat—an area projected to contract under continued regional warming [15]—the SG fishery is particularly vulnerable to climate-driven shifts.

These external shocks contribute to the high-risk nature of krill fisheries in this region [9].

Our study demonstrates that krill recruitment to SG follows dominant transport pathways, which in turn influence fishing patterns in the following season. We find that AP serves as the primary source region for SG krill, with interannual variability in AP transport affecting recruitment strength and spatial distribution. In contrast, South Orkney (SO) is a secondary source, with weaker direct influence on SG fisheries, though some connectivity exists between the two regions.

These findings underscore the need for dynamic fisheries management that accounts for environmental variability and changing transport dynamics. By integrating simulation-based insights into fisheries planning, stakeholders can better anticipate fluctuations in krill availability and adapt management strategies to ensure long-term sustainability in the Southern Ocean.

## Acknowledgments

Project funding was provided through the Research Council of Norway as part of the SFI Harvest project (986010103).

